# Explosive shock wave exposure leads to age-accelerated motor and sensory decline in *C. elegans*

**DOI:** 10.64898/2025.12.19.695190

**Authors:** Jessica Tittelmeier, Henrik Seeber, Dennis Grasse, Nirav Barapatre, Christoph Schmitz, Daniel Krentel, Steffen Grobert, Carmen Nussbaum-Krammer

## Abstract

**Background:** Military and law enforcement personnel are routinely exposed to shock waves from weapon systems and explosive devices during training and operations. Even when such shock wave exposure results in mild traumatic brain injury (mTBI) without abnormalities on routine imaging, persistent neurological symptoms may occur. Yet, the biological processes that link an acute blast wave insult to progressive neuronal dysfunction remain poorly defined.

**Methods:** We established *Caenorhabditis elegans* nematodes as a genetically accessible animal model for blast-related mTBI (br-mTBI). Animals were exposed in gelatin to shock waves generated by a custom-built shock wave generator (SWG), which produces explosive shock waves with an abrupt overpressure peak followed by a negative pressure phase. The resulting pressure-time curves are very similar to the profiles of conventional explosives in free-field setups. To enable mechanistic studies under controlled laboratory conditions, we developed a complementary platform using a medical radial extracorporeal shock wave therapy (rESWT) device. Motor behavior, mechanosensory function, neuronal morphology and cytoskeletal integrity were analyzed during aging.

**Results:** SWG-derived shock waves induced immediate but reversible motor and sensory deficits, followed by an accelerated age-dependent decline in movement and touch sensitivity. The rESWT platform accurately reproduced these phenotypes. Touch receptor neurons showed progressive structural abnormalities, including degeneration, alongside acute PTL-1/Tau mislocalization consistent with cytoskeletal injury.

**Conclusions:** Defined shock wave exposure is sufficient to provoke long-term neuronal functional decline in *C. elegans*, accompanied by structural deterioration and premature neurodegeneration. This tractable model enables lifelong phenotyping and mechanistic dissection of how an acute shock wave insult progresses to chronic neuronal dysfunction, and provides a scalable platform for identifying molecular and pharmacological modifiers that promote resilience.

## BACKGROUND

Blast waves from heavy weapons and explosives pose a significant risk of traumatic brain injury (TBI) to military personnel and specialized police units. During routine breaching or weapon training, personnel can experience repeated low-level blast (LLB) exposure that may not cause visible structural damage but can still produce long-lasting neurological and cognitive symptoms [1–3]. These mild or sub-concussive TBIs often go undetected by conventional clinical imaging, yet they are increasingly linked to chronic impairments and higher risk of dementia later in life [1–12].

Explosive shock waves are characterized by a steep positive pressure rise, high peak overpressure, a subsequent negative-pressure phase and a short duration with rapid decay [13, 14]. As the shock wave propagates through the head, it encounters structures that vary in density, composition and characteristics, resulting in heterogeneous transmission of energy. At interfaces with differing acoustic impedance – such as between the skull, brain parenchyma and cerebrospinal fluid – reflection and refraction of the shock wave front further lead to local energy accumulation and increased interface damage [4, 15, 16].

At high intensities, shock waves can induce transient cranial deformation and intracranial pressure changes resembling blunt-impact injuries [17, 18]. In contrast, LLBs, as typically experienced during training, do not cause any gross structural lesions or hemorrhages [17, 18]. Instead, microscopic damage such as axonal, synaptic and endothelial injuries without prominent tissue disruption predominates. In humans, diffusion tensor MRI shows diffuse white-matter abnormalities despite normal clinical findings [11, 19], and post mortem analyses reveal astroglial scarring at perivascular and gray–white matter interfaces [15]. Corresponding animal models demonstrated microvascular dysfunction, blood-brain barrier disruption, axonal and synaptic damage, mitochondrial dysfunction and early neuroinflammatory responses following low-intensity shock wave exposure – despite the absence of macroscopic pathology [16, 20–23].

The biological processes linking the transient mechanical insult of a shock wave to delayed neuronal dysfunction have remained poorly understood. Elucidating these mechanisms is crucial for developing therapeutic interventions and protective measures. However, the investigation of secondary and chronic effects of blast-related mild traumatic brain injury (br-mTBI) in mammalian models is constrained by ethical considerations, high costs, low experimental throughput, considerable interindividual variability and the difficulty to expose transgenic animals to shock waves outside of shielded facilities – all factors that impede progress in elucidating fundamental mechanisms [24].

The aim of this study was to establish the small invertebrate nematode *Caenorhabditis elegans* (*C. elegans*) as a tractable system to accelerate research on br-mTBI. *C. elegans* has become one of the most widely used and versatile model organisms in contemporary biological and genomic research, owing to its unique combination of genetic accessibility, cellular resolution and lifetime observability [25]. The complete identity, morphology and synaptic connectivity of all 302 neurons are known, and its optical transparency enables longitudinal imaging of neuronal structure and function throughout the lifespan [26–29]. Combined with a powerful genetic toolkit and short lifespan, these features enable rapid analysis of conserved molecular and metabolic pathways during aging [26, 27, 30, 31]. In contrast to mammalian models such as mice, *C. elegans* provides direct experimental access to individual neurons and complete neural circuits, enabling the correlation of cellular changes with complex behavioral readouts [26–29]. Because *C. elegans* has no centralized brain, blood–brain barrier or vasculature, it enables clean isolation of neuron-intrinsic responses to shock wave exposure. These characteristics make *C. elegans* ideally suited for investigating neuronal injury and stress responses at both cellular and systems levels.

*C. elegans* has been used to model diverse neurotoxic and neurodegenerative processes, including the toxicity of disease-associated protein aggregation or chemical warfare agents such as organophosphates [32–35]. Of note, a *C. elegans* model of blunt force trauma revealed conserved metabolic mechanisms that also protect mammalian neurons [36], underscoring its translational potential.

Here, a custom-built shock wave generator (SWG) was employed that produces explosive shock waves under free-field-like conditions [13, 14, 37]. This system is based on the controlled detonation of an acetylene-oxygen mixture in a cylindrical autoclave, releasing shock waves through a defined outlet [13, 14]. While the overall propagation of the fluid is not perfectly spherical due to the directed outlet, the shock waves expand quasi-hemispherically and undisturbed within the angle of interest next to the front of the autoclave, where biological and material samples can be positioned [13, 14]. In this region, the pressure-time profiles are characterized by a steep rise, peak overpressures of approximately 80 ± 10 kPa at 1 m distance, and an exponential decay followed by a negative phase, closely resembling ideal blast waveforms with additional weapon system muzzle blast-like features. Compared to conventional explosive-based testing or shock tubes, the SWG offers safe, scalable, undisturbed, highly reproducible and physiologically relevant LLB conditions for both material and biological research [13, 14], including short turn-around phases to enable a high test frequency.

To establish a *C. elegans* model of br-TBI under these conditions, nematodes were pipetted in liquid into cavities within gelatin blocks that replicate the transmission and attenuation of shock waves through soft tissue such as the brain, with specific acoustic characteristics (e. g. sound-propagation velocity of ~1550 m/s compared to ~340 m/s in air [38]). In a second step, a laboratory shock wave model was implemented using a medical radial extracorporeal shock wave therapy (rESWT) device. This setup enabled replication of the shock wave-induced phenotypes under defined laboratory conditions suitable for mechanistic studies. Animals exposed to radial extracorporeal shock waves (rESWs) showed immediate motor impairments with transient recovery, followed by an age-accelerated decline in movement and mechanosensory function. Touch receptor neurons (TRNs) developed progressive morphological abnormalities and degeneration, and an acute mislocalization of the Tau homolog PTL-1 from neurites to somata, consistent with cytoskeletal damage. Thus, this model recapitulates key aspects of br-TBI and provides a platform to dissect the molecular mechanisms through which a transient shock wave insult evolves into delayed motor and mechanosensory decline in *C. elegans*.

## METHODS

### Shock Wave Generator setup

Free-field experiments were carried out at the German Federal Institute for Materials Research and Testing (Bundesanstalt fuer Materialforschung und -pruefung, BAM) Test Site for Technical Safety (Testgelaende Technische Sicherheit, TTS) in Baruth/Mark (Germany) in cooperation with the Bundeswehr (German Federal Armed Forces). The experiments used a SWG based on a cylindrical autoclave as described in [14] (Figure 1A-E).

**Figure 1.**
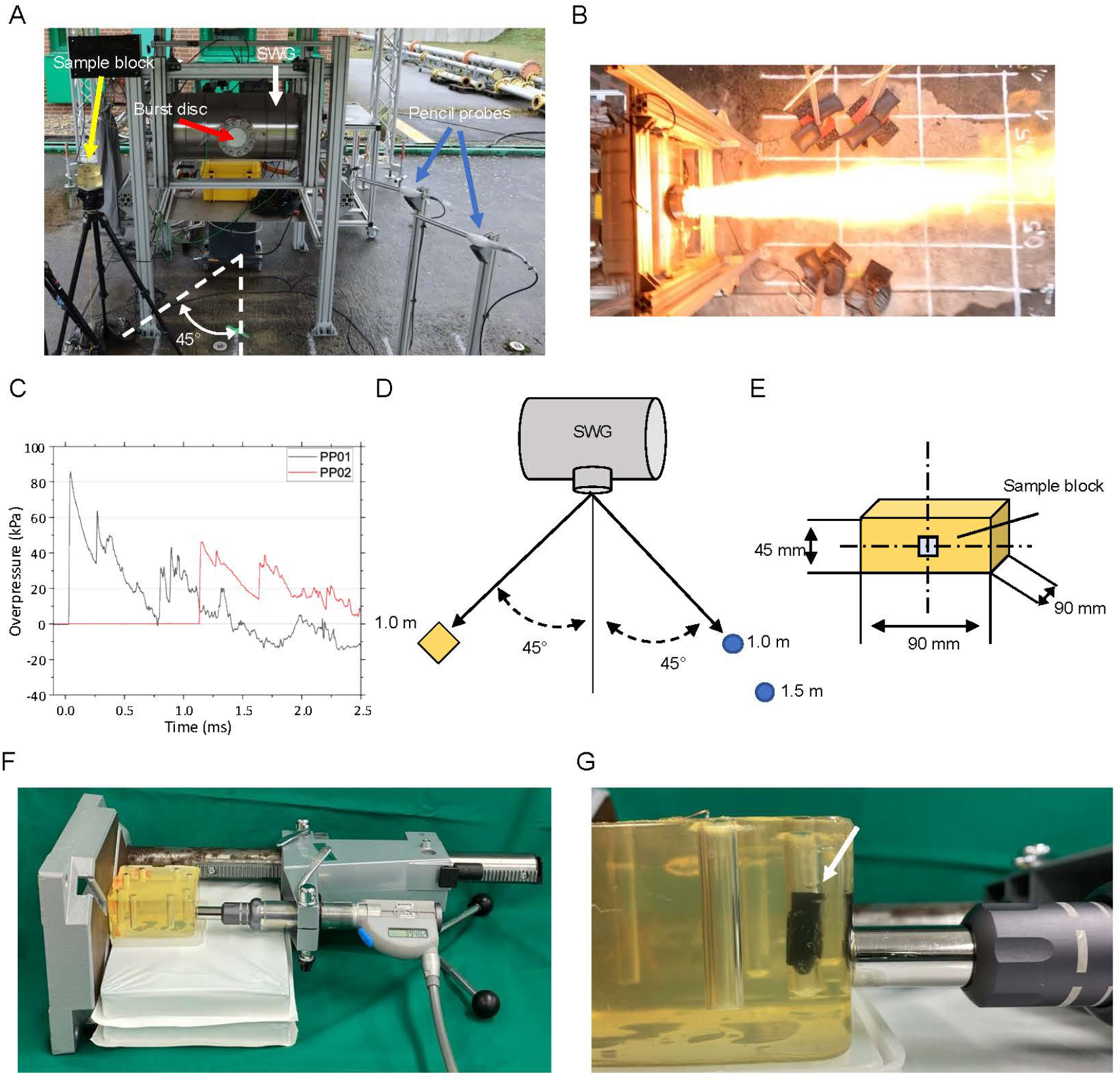
Experimental configurations for free-field shock wave generation and laboratory-based radial extracorporeal shock wave (rESW) delivery. (**A**) Photograph of the free-field setup showing the shock wave generator (SWG, white arrow), the explosion outlet sealed with an aluminum burst disc (red arrow), a gelatin block positioned 1.0 m from the SWG (yellow arrow) and the pencil pressure probes (blue arrows). (**B**) Photograph capturing the SWG explosion. (**C**) Representative pressure–time curves recorded by pencil probes placed 1.0 m (black curve) and 1.5 m (red curve) from the SWG, showing an initial overpressure peak characteristic of an ideal blast waveform. (**D**) Schematic illustration of the free-field arrangement. Gelatin blocks (yellow) were positioned 1.0 m from the SWG within a 45° angle, and pencil probes (blue) were placed at 1.0 m and 1.5 m. (**E**) Schematic side view of the gelatin block containing *C. elegans* samples (light blue square). (**F**) Overview of the laboratory setup for exposing *C. elegans* to radial extracorporeal shock waves (rESWs). The handpiece of the medical rESWT device was held in a fixed position using a drill stand to ensure consistent delivery of rESWs. (**G**) Close-up view of the rESW applicator in contact with the gelatin block, showing a black gelatin capsule inserted into a cavity within the block (white arrow).

The SWG consisted of a 65 l (0.065 m^3^) autoclave (white arrow in Figure 1A) filled with a homogenous, stoichiometric acetylene–oxygen mixture under atmospheric pressure. Ignition was initiated with an electric wire inside the vessel, causing a gas detonation, the jet of fire (Figure 1B) and the shock wave to exit through a 12.5 cm diameter outlet sealed by an aluminum burst disc (red arrow in Figure 1A). The outlet center was positioned 1.35 m above the ground, and the resulting shock wave propagated quasi-hemispherically and undisturbed at a distance of 1.0 – 1.5 m from the outlet (Figure 1C) [13, 14]. The sample block (yellow arrow in Figure 1A, yellow block in Figure 1D, E) and pencil probes (blue arrows in Figure 1A and blue circles in Figure 1D; Type 6233AA0025; Kistler Instruments, Winterthur, Switzerland; pressure sensors used to characterize the shock wave and thus the load case) were placed at 45° (indicated in Figure 1A, D), thus avoiding the jet of fire.

### Gelatin blocks

Gelatin (#63043, Kremer Pigmente, Aichstetten, Germany) was dissolved in H_2_O to obtain a 17% gelatin (w/v) solution and allowed to absorb H_2_O for 2 h at room temperature. The solution was then placed in a 55 °C water bath and slowly heated up to this temperature until it was clear, and subsequently poured into a block mold. Plastic rods were inserted 1 cm from the edge to form cavities and removed once the gelatin had solidified. A 17% gelatin concentration was chosen because it exhibits a sound-propagation velocity of 1542.42 m/s at 15 °C, which closely resembles that of human soft tissue [39]. Gelatin capsules were inserted into cavities within the gelatin block, and *C. elegans* samples were then pipetted in standard M9 low salt buffer (21 mM Na_2_HPO_4_, 22 mM 1KH_2_PO_4_, 85 mM NaCl, 1 mM MgSO_4_) into gelatin capsules before exposure (Figure 1E).

### rESWT device setup

Radial extracorporeal shock waves were generated using a medical rESWT device (Swiss DolorClast; Electro Medical Systems, Nyon, Switzerland). This device produces rESWs ballistically by accelerating a projectile through compressed air. The projectile then strikes a metal applicator, converting kinetic energy into a radially expanding pressure wave [40–43]. For all experiments, the device was operated at 2 bar air pressure and 5 Hz frequency. The applicator was positioned directly against the gelatin block to deliver consistent exposure (Figure 1F, G).

### Maintenance of *C. elegans* and age synchronization

*C. elegans* were cultured using standard methods [44]. Animals were grown on nematode growth medium (NGM) plates seeded with *E. coli* strain OP50 at 20 °C. The strains used in this study included wild-type N2 as well as the transgenic strains CNK166 (mec-4p::mCherry) and PTL-1::mNeonGreen [34, 45]. The strain CNK166 (mec-4p::mCherry) expresses mCherry under the control of the TRN-specific mec-4 promoter, enabling visualization of TRN somata and neurites. PTL-1::mNeonGreen transgenic animals express mNeonGreen-tagged PTL-1, the *C. elegans* homolog of human microtubule-associated protein Tau (MAPT/Tau), under its own promoter, which is predominantly active in TRNs [46]. This strain was used to examine microtubule integrity.

Animals were age-synchronized by bleaching. Briefly, gravid adults were dissolved in 20% sodium hypochlorite solution. The surviving *C. elegans* embryos were hatched overnight in M9 buffer with gentle rocking at 20 °C. The next day, appropriate amounts of L1 larvae were added to the plates.

### Motility assay

Age-synchronized *C. elegans* were transferred into M9 buffer from NGM plates and were given time to adapt to the new environment. The number of body bends within 30 s was counted to determine the swimming speed. One thrash was defined as the entire motion from one side to the other. 10 animals were examined for each condition in 2-3 biological replicates.

### Mechanosensory assay

Mechanosensory assay was done as described in [47]. Specifically, gentle touch sensitivity was assessed by brushing an eyebrow hair (attached to a toothpick) across either the anterior region of *C. elegans*, just posterior to the pharynx, or the posterior region. Ten animals were examined on each day for each condition in 2-3 biological replicates with 5 strokes each. A positive touch response was counted when the animal stopped or moved away from the stimulus.

### Mounting of live *C. elegans* for imaging

Age-synchronized animals were mounted on 8-10% agarose in M9 buffer pads with a drop of mounting mix (2 mM levamisole and 50% (v/v) nanosphere size standards (Thermo Fisher, Waltham, MA, USA)) and covered with a coverslip.

### Assessment of touch receptor neuron morphology

The strain CNK166 was used for neurotoxicity scoring. For consistency, we focused on the posterior touch receptor neurons (PLMs), as they are easily visualized in the posterior region of the animal, and their morphology can be reliably assessed using the mCherry signal. Live animals were imaged using an Olympus Ixplore SpinSR Confocal microscope with an UplanS Apo 60x/1.30 Silicon oil objective (Olympus, Tokyo, Japan). The neurotoxicity score was determined by adding a score of 1 for each of the following phenotypes: soma outgrowth, process branching, and 3 points for degeneration, when the soma was no longer visible. 10 animals were examined at each day of age in 3 independent biological replicates.

### Assessment of PTL-1::mNeonGreen distribution

To quantify the relative distribution of PTL-1::mNeonGreen between the soma and the neurites of the anterior touch receptor (ALM) neuron, where PTL-1 is highly expressed, confocal images were acquired using the confocal microscope described above (Olympus iXplore SpinSR). All images were obtained under identical acquisition settings, 2 h post-exposure. Imaging the anterior region of the animals enabled simultaneous analysis of the ALM neuron and the surrounding hypodermis. For the soma/neurite ratio, a median focal plane (z-plane) passing through the center of the soma was selected. Two regions of interest (ROIs) were manually drawn using Fiji (ImageJ) [48], one encompassing the neuronal soma and another along the proximal neurite, immediately distal to the soma. The mean fluorescence intensity within each ROI was measured, and the soma/neurite ratio was calculated as the mean intensity of the soma divided by that of the neurite.

To quantify hypodermal PTL-1::mNeonGreen, an ROI was drawn around the hypodermis. Within this ROI, the Moments threshold mask in Fiji was applied to identify fluorescent signal. The integrated density (IntDen in Fiji) was obtained to derive a combined measure of hypodermal PTL-1::mNeonGreen fluorescence.

### Statistical analysis

Statistical analysis was performed with GraphPad Prism (Version 5.04; GraphPad Software, Boston, MA, USA). Data was tested for normal distribution by D’Agostino and Pearson omnibus normality test. Data presentation, sample size (number of biological repeats and sample size per repeat and condition) and the applied statistical tests are indicated in the figure legends for each experiment. Significance levels: non-significant (n.s.) p > 0.05, *p ⩽ 0.05, **p ⩽ 0.01 and ***p ⩽ 0.001.

## RESULTS

### Shock wave exposure triggered acute, reversible motor deficits and accelerated age-dependent motor and sensory decline in *C. elegans*

To investigate the biological consequences of shock wave exposure in *C. elegans*, we adapted the SWG setup [13, 14] to a tissue-relevant configuration [37]. *C. elegans* pipetted in liquid into cavities within gelatin blocks were exposed to shock waves produced by the SWG at 1 m distance and 45° from the outlet [13, 14] (Figure 1A). This setup exposed the animals to the mechanical load of the shock wave while avoiding confounding influences from the jet of fire, exhaust gases or flow artifacts (Figure 1B). Representative pressure-time curves recorded from pencil probes at 1.0 m and 1.5 m distance confirmed characteristic shock waves with a steep initial rise and a short negative phase (Figure 1C). The peak overpressure in the free-field was 88.3 kPa (1.0 m) and 46.3 kPa (1.5 m), respectively, similar to previous studies using this setup [13, 14].

*C. elegans* populations were exposed to one single shock wave with the characteristics described above on the first day of adulthood (A1, corresponding to day 4 of life) and subsequently transferred from the cavities within the gelatin block back to standard NGM plates for longitudinal analysis at defined time points during aging. Control animals underwent the same handling procedure (including pipetting into gelatin cavities, remaining in the blocks for the same duration, and transfer back to NGM plates), but their gelatin blocks were placed outside the SWG exposure zone at the back of the building, ensuring they experienced all manipulations except the shock wave itself.

One hour after exposure, animals exhibited a marked reduction in motor function (Figure 2A). By the next day (A2, 24 h post-exposure), motor performance had returned to control levels, indicating that the acute impairment was reversible. From day 3 of adulthood (A3, two days post-exposure) onward, however, shock wave-exposed *C. elegans* exhibited an accelerated, age-dependent decline in motor function relative to controls (Figure 2B). Thus, shock wave exposure triggered an acute but reversible motor deficit, followed by an accelerated decline in motor function during aging.

**Figure 2.**
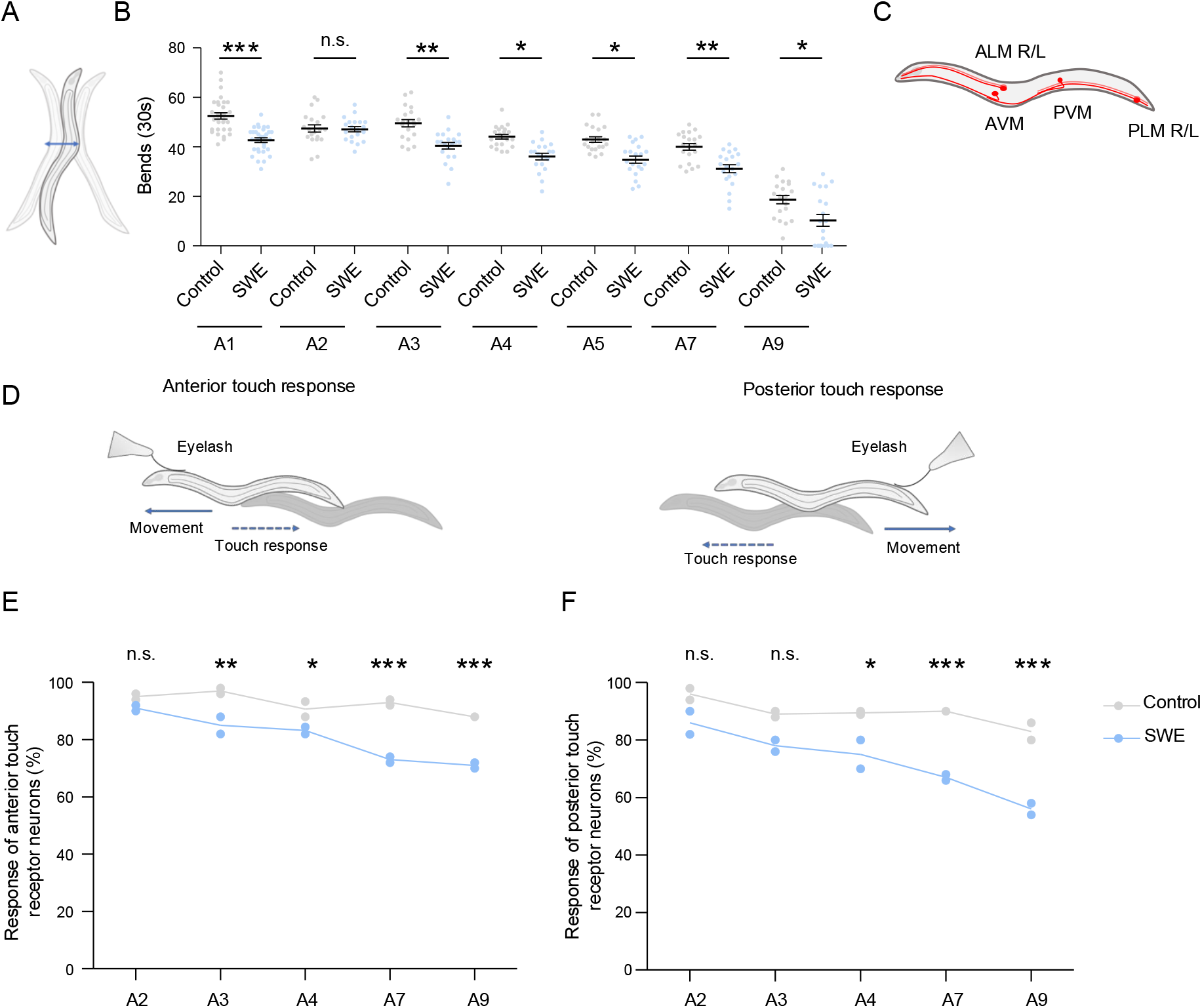
Shock wave exposure induced acute motility impairments and accelerated touch receptor neuron (TRN) functional decline. (**A**) Schematic representation of *C. elegans* swimming movements (**B**) Quantification of body bends during a 30-s swimming period at the indicated ages. Compared to control animals (Control), shock wave-exposed animals (SWE) showed an acute reduction in motility at A1 (1 h post-exposure), followed by an accelerated decline during aging (A3–A9). Statistical significance was assessed using two-way ANOVA with Bonferroni correction. Each dot represents one animal; data were obtained from two independent biological replicates (10 animals per replicate). Lines indicate mean ± SEM. (**C**) Schematic of the six touch receptor neurons (TRNs: ALM R/L, AVM, PVM, PLM R/L). (**D**) Schematic of the touch response assay used to assess anterior and posterior TRN function. (**E, F**) Quantification of anterior (E) and posterior (F) TRN responses to gentle touch across adulthood (A2–A9). TRN function was comparable to control animals (Control) in early adulthood (A2–A3) but declined more rapidly from A4 onward in shock wave-exposed animals (SWE). Statistical significance was determined using two-way ANOVA with Bonferroni correction. Data points represent biological replicates (10 animals per replicate); lines indicate means. *p < 0.05, **p < 0.01, ***p < 0.001, n.s. = not significant.

The TRNs are specialized mechanosensory neurons that mediate gentle touch responses through well-characterized neural circuitry [47]. Because they respond directly to mechanical stimuli, TRNs may be particularly vulnerable to shock waves. In *C. elegans*, touch sensation is mediated by a defined set of six TRNs: the anterior neurons ALML, ALMR and AVM, and the posterior neurons PLML, PLMR and PVM (Figure 2C). Gentle touch to the anterior or posterior part of the animals triggers a stereotyped withdrawal response controlled by these neurons [34, 47] (Figure 2D). To determine whether TRN function was affected by shock wave exposure, we quantified the anterior and posterior touch response across aging (Figure 2E, F). In early adulthood, TRN function was indistinguishable between shock wave-exposed and control animals: anterior touch responses remained unchanged at A2 and posterior responses remained unchanged from A2 to A3 (Figure 2E, F). At later ages, however, both anterior and posterior touch responses declined more rapidly in shock wave-exposed nematodes (Figure 2E, F), indicating an accelerated loss of mechanosensory function. Logistical and time constraints at the free-field testing site prevented assessment of acute effects within the first 24 h post-exposure. A transient early impairment with subsequent recovery, similar to motor function, therefore remains possible. Nevertheless, TRN function was unaffected in young adults (A2, 24h post-exposure) but declined more rapidly thereafter, consistent with a shock wave-induced acceleration of age-dependent mechanosensory deterioration.

### A controlled rESWT platform bypassed free-field constraints of the SWG and recapitulated acute and age-accelerated neurobehavioral phenotypes in *C. elegans*

Although the SWG represents a major advance over conventional explosives, its mandatory free-field setup still presents significant challenges that impede progress in uncovering the molecular mechanisms of shock wave– induced neuronal injury in *C. elegans*. Specifically, transgenic animals could not be exposed to shock waves under free-field conditions, and the setup prevented the use of high-end microscopes for immediate post-exposure behavioral assessment. Additional practical limitations included the need to transport mobile laboratory equipment such as incubators.

To overcome these limitations, we implemented a laboratory-based method using a well-characterized medical rESWT device (Swiss DolorClast; Electro Medical Systems) (Figure 1F, G) [40, 42, 49, 50]. In this device, compressed air accelerates a projectile within a guiding tube, and its impact on a metal applicator produces a radially expanding pressure wave. The resulting rESWs display hallmark characteristics of shock waves in biological tissue, such as a steep pressure rise, high peak amplitude and a brief tensile phase [40, 50]. At 2 bar driving pressure and 5 Hz repetition rate, this device produces rESWs with a mean peak positive pressure of approximately 2.5 MPa at a distance of 10 mm from the applicator (measured in water), with a positive phase lasting ~3-4 μs [49].

Although these rESWs share the rapid pressure rise and short negative pressure phase characteristic of shock waves, their peak positive pressure is different to the SWG-derived shock waves. We therefore empirically tested repetitive rESWs (100, 200, 300, 500) to identify conditions that best reproduced the SWG phenotypes. Unlike earlier studies in which *C. elegans* were exposed in liquid medium in a 96-well format [41, 51], nematodes were exposed through gelatin, analogous to the SWG setup, to ensure comparable rESW propagation. Animals exposed to increasing numbers of rESWs showed an initial decline in motility, similar to animals exposed to SWG-derived shock waves (Figure 3A). However, nematodes exposed to 500 rESWs did not recover to control levels by A2, unlike those exposed to a single SWG-derived shock wave (Figure 3A). We therefore selected 300 rESWs for detailed analysis, since animals showed a clear impairment at A1 that was fully reversible at A2 (Figure 3A). Under this condition, nematodes displayed the same trajectory as observed with the SWG setup: an acute decline in motility, transient recovery and an accelerated age-related decline (Figure 3B).

**Figure 3.**
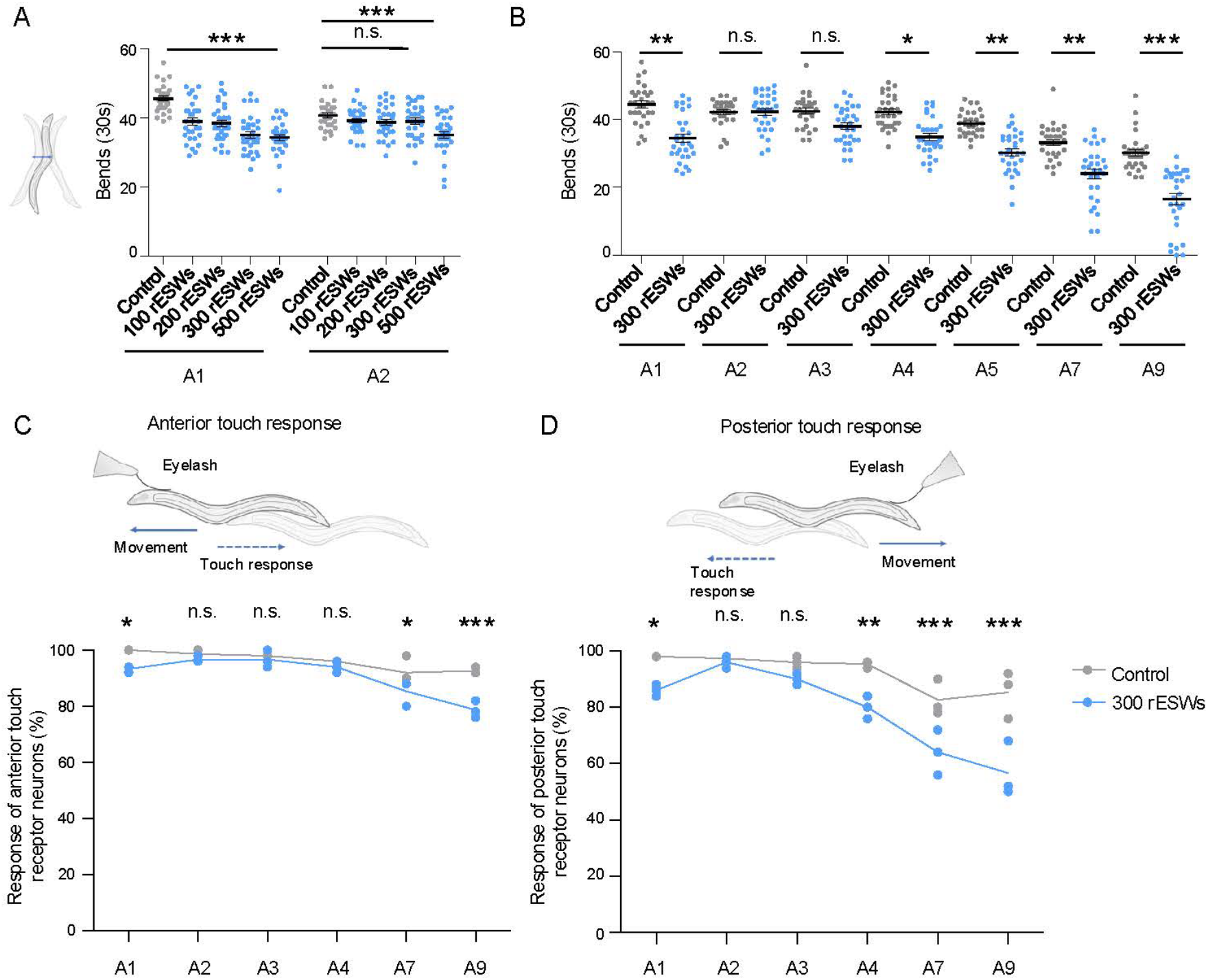
Exposure to 300 rESWs phenocopied motility decline and touch receptor neuron (TRN) dysfunction after SWG-derived shock wave exposure. (**A**) Quantification of body bends during a 30-s swimming period of *C. elegans* exposed to 100–500 radial extracorporeal shock waves (rESWs) at 2 bar and 5 Hz, measured 1 h (A1) and 24 h (A2) post-exposure. All conditions showed an acute motility decline at A1, with recovery at A2 except at 500 rESWs. Statistical significance was assessed using a Kruskal–Wallis test with Dunn’s comparison. Each dot represents one animal; data are from three independent biological replicates (10 animals per replicate). Lines indicate mean ± SEM. (**B**) Quantification of body bends during a 30-s swimming period of *C. elegans* exposed to 300 rESWs at the indicated ages. Animals displayed an acute motility reduction at A1 with recovery by A2, followed by an accelerated decline during aging (A4–A9). Statistical analysis was performed using two-way ANOVA with Bonferroni correction. Each dot represents one animal; data are from three independent biological replicates (10 animals per replicate). Lines indicate mean ± SEM. (**C, D**) Quantification of TRN responses to gentle touch across adulthood after exposure to 300 rESWs. TRN function showed an initial decline 1 h post-exposure, no significant differences from controls at early adult stages (A2–A3), and significantly reduced responses emerging from A7 (anterior; C) or A4 (posterior; D) onward in rESW-exposed nematodes. Statistical significance was determined using two-way ANOVA with Bonferroni correction. Data points represent biological replicates (10 animals per replicate); lines indicate means. *p < 0.05, **p < 0.01, ***p < 0.001, n.s. = not significant.

The TRN function showed a similar pattern. 1h post-exposure, both anterior and posterior TRNs exhibited reduced mechanosensory responses (Figure 3C, D). Responses returned to control levels at A2–A4 (anterior) and A2–A3 (posterior), but declined more rapidly thereafter, paralleling the progressive sensory deficits observed with the SWG setup (Figure 3C, D). Thus, repetitive exposure to rESWs accurately reproduced the acute and long-term behavioral and neuronal phenotypes observed after SWG-derived shock wave exposure.

### Accelerated TRN degeneration and PTL-1/Tau mislocalization revealed structural mechanisms of rESW– induced sensory impairment

To determine whether sensory deficits reflected structural neuronal damage, we examined TRN morphology using transgenic animals expressing mCherry under the mec-4 promoter (mec-4p::mCherry) [34] (Figure 4A). Posterior TRNs (PLM) displayed characteristic age-associated abnormalities, including soma outgrowths, ectopic branching and degeneration, which increased with age in controls [34, 52] but were significantly more frequent after exposure to 300 rESWs (Figure 4A, B). These findings indicate that the functional decline of TRNs was accompanied by an accelerated accumulation of morphological abnormalities during aging.

**Figure 4.**
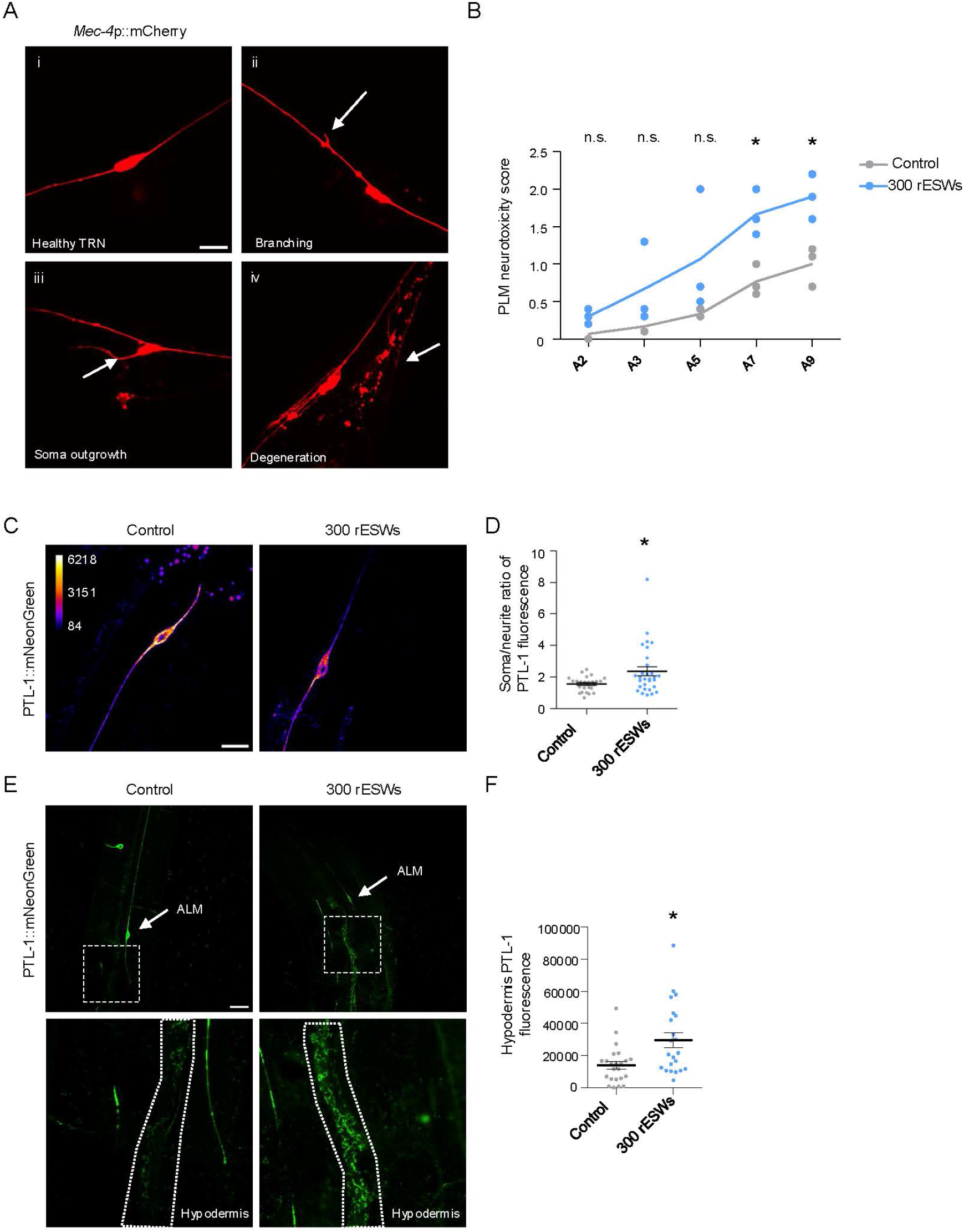
Exposure to 300 rESWs induced structural and molecular damage in touch receptor neurons (TRNs). (**A**) Representative examples of in vivo confocal images of TRNs expressing mec-4p::mCherry, illustrating normal morphology (i), branching (ii), soma outgrowth (iii) and degeneration (iv). (**B**) Quantification of PLM neurotoxic phenotypes showed an age-dependent increase in morphological abnormalities, which was significantly accelerated after exposure to 300 rESWs. Statistical significance was assessed using two-way ANOVA with Bonferroni correction. Data points represent biological replicates (10 animals per replicate); lines indicate means. (**C**) Representative examples of in vivo confocal images of ALM neurons expressing PTL-1::mNeonGreen (shown with an intensity LUT). (**D**) Quantification of the ALM soma/axon fluorescence ratio showed reduced axonal PTL-1 signal after exposure to 300 rESWs. Statistical analysis was performed using a Mann–Whitney test. Each dot represents one animal; data are from three independent biological replicates (10 animals per replicate). Lines indicate mean ± SEM. (**E**) Representative examples of in vivo confocal images of *C. elegans* expressing PTL-1::mNeonGreen. (**F**) Quantification of hypodermal PTL-1::mNeonGreen fluorescence, revealing increased PTL-1 accumulation after exposure to 300 rESWs, consistent with enhanced protein redistribution. Statistical analysis was performed using a Mann–Whitney test. Data are presented as mean ± SEM. *p < 0.05, n.s. = not significant. Scale bar: 10 μm.

Because diffuse axonal damage has been observed in br-mTBI patients and vertebrate models [53–55], we examined neurite integrity using *C. elegans* expressing PTL-1::mNeonGreen, the homolog of human MAPT (Tau) [27, 45]. PTL-1 serves as a sensitive marker of microtubule stability and axonal health. Following exposure to 300 rESWs, animals exhibited reduced PTL-1 fluorescence in the neurite (Figure 4C, D), indicative of PTL-1 detachment from neurites.

In parallel, we observed increased PTL-1 accumulation in the surrounding hypodermis (Figure 4E, F). In *C. elegans*, TRNs are embedded within the hypodermis, and neuronal debris has been shown to undergo transcellular degradation in this tissue [34, 56, 57]. The accumulation of PTL-1 in the hypodermis after exposure to rESWs likely reflects the transfer of damaged PTL-1 for degradation. Together, these findings demonstrate that rESWs cause structural as well as molecular damage.

This model provides a powerful framework to dissect how acute blast injury translates into lasting neuronal dysfunction.

## DISCUSSION

This study introduces *C. elegans* as a scalable and genetically tractable model for br-mTBI. By combining free-field-based experiments using a SWG with a laboratory setup employing a medical rESWT device, we created a complementary platform to dissect acute and long-term molecular and cellular responses to shock wave-induced injury.

In both the free-field SWG and the laboratory rESW setup, *C. elegans* exhibited an acute reduction in motility after exposure that fully recovered, but was followed by a markedly accelerated decline in motor and sensory function with age. This temporal pattern is consistent with human data showing that shock wave exposure – even when acute symptoms (e.g., loss of consciousness, dizziness) were not recognized [58] – can be followed by delayed cognitive, affective and degenerative outcomes over weeks to years [7, 59]. The early phase likely reflects direct mechanical disturbance of neuronal integrity, whereas the delayed phase may involve secondary cellular stress pathways. Dissecting these phases in *C. elegans* could identify molecular mechanisms that determine neuronal vulnerability and resilience after blast exposure.

In our previous studies, *C. elegans* were exposed in liquid-filled 96-well plates, which caused severe paralysis, only partial recovery and shortened lifespan even at a relatively low dose of 100 rESWs [41, 42, 51]. These pronounced effects may reflect forces beyond the primary pressure pulse: the tiny wells may have caused pressure reflections, the immersed applicator may have caused rapid, turbulent flows that swirled the animals around, and the confined space may have made collisions with the walls likely, adding a component of blunt force impact. In the current setup, animals were exposed through gelatin blocks with >1 cm standoff from the applicator, so exposure was dominated by a propagating rESW rather than turbulence or contact forces. This refinement produced a milder yet persistent phenotype and more closely models injury by the primary shock wave.

Furthermore, in both the free-field SWG and the laboratory rESW setup, TRNs exhibited an accelerated functional decline after exposure. These neurons are directly coupled to the cuticle and rely on specialized microtubule bundles that are tethered to membrane through proteins such as MEC-2 [47]. This architecture enables high mechanosensitivity but also renders TRNs particularly vulnerable to mechanical overload. Shock wave-induced perturbations in membrane tension and microtubule stability are consistent with axonal injury mechanisms observed in patients and vertebrate br-mTBI models [53–55]. Our findings suggest that mechanical stress triggers conserved pathways of microtubule destabilization, which may represent early events in trauma-induced neurodegeneration.

We deliberately chose not to introduce a skull-like interface between the shock wave source and the animals. Such a structure would attenuate the shock wave substantially – transmitting only about 16% of the energy [60] – and introduce strong heterogeneity in exposure depending on each animal’s position and orientation relative to the skull surface. Moreover, the biomechanical scaling between *C. elegans* and the human brain is not directly translatable. Instead, our focus was to resolve microscopic neuronal damage and track neuronal fate and function during aging after a defined shock wave-induced insult. This approach enables a high-resolution, genetically accessible analysis of how this acute physical trauma evolves into chronic proteotoxic and neurodegenerative changes.

A limitation of this study is that the pressure profiles generated by the SWG in the free-field setup cannot be directly compared with those produced by the rESWT device in the laboratory. For this reason, the laboratory model was optimized empirically to reproduce the behavioral phenotypes seen after SWG-derived shock wave exposure, rather than to match its exact physical parameters. This approach, however, offers several advantages. First, the rESW-based platform supports much higher experimental throughput at lower cost and can be run entirely under standard laboratory conditions, enabling rapid iteration, tighter control of confounding variables and detailed longitudinal phenotyping. Second, calibrating the laboratory exposure to recapitulate the SWG-induced motor and sensory trajectories emphasizes construct validity while avoiding the logistical constraints of free-field experiments. Finally, the aim of establishing the laboratory model was not to replicate free-field conditions exactly, but to provide a tractable, well-controlled system for dissecting the molecular and cellular mechanisms of br-mTBI. This platform enables genetic and pharmacological manipulation, high-resolution imaging and stress response analyses, with the goal of identifying mechanistic biomarkers and candidate modifiers that can later be validated under free-field SWG conditions.

## Conclusion

Exposure to a defined shock wave resulting from a gas detonation was sufficient to induce immediate functional deficits in *C. elegans*, followed by transient recovery and subsequent progressive structural and functional deterioration of vulnerable neurons. This trajectory mirrors clinically relevant patterns in which early recovery after LLB exposure can mask the emergence of delayed neurological decline.

By integrating free-field SWG experiments with a complementary laboratory rESW setup, this study established a powerful platform for dissecting shock wave–induced injury processes with cellular and molecular resolution. The *C. elegans* model provides sensitive tools for detecting subclinical early molecular and structural damage, tracking its progression over time and linking these early events to long-term functional impairment. This approach bridges the gap between complex mammalian br-mTBI models and reductionist cellular assays.

Mechanistic insights gained from this system may reveal molecular and genetic modifiers that promote neuronal resilience after blast injury. These candidates can then be specifically validated in mammalian models and human cohorts, accelerating the identification of therapeutic targets and development of protective strategies – an urgent need given the ongoing occupational risk faced by military and law enforcement personnel.

## STATEMENTS AND DECLARATIONS

## Acknowledgements

We thank Sabine Tost for excellent technical assistance. The PTL-1::mNG strain was generously provided by Dr. Miriam B. Goodman (Stanford University). Some strains were provided by the Caenorhabditis Genetics Center (CGC), which is funded by NIH Office of Research Infrastructure Programs (P40 OD010440).

## Availability of data and materials

The datasets used and analyzed during this study are available from the corresponding author on reasonable request.

## Ethics approval

Not applicable.

